# The Impact of Germ Layer Origin on the Ageing Transcriptome

**DOI:** 10.64898/2026.11.20.768114

**Authors:** Tamsin Ederies, João Pedro de Magalhães

## Abstract

Ageing is accompanied by widespread changes in gene expression, yet whether these changes share any common organisation across groups of tissues remains unclear. Adult tissues arise from three embryonic germ layers (ectoderm, mesoderm and endoderm), so shared developmental lineage could influence how tissues age. Using bulk RNA-seq data from 24 GTEx tissues, we estimated age effects in each tissue with limma-voom and combined them within each germ layer by random-effects meta-analysis. Endoderm had the most significant age-associated genes (4,805), followed by mesoderm (4,520) and ectoderm (1,744), but this order reversed for genes with consistent effects across tissues of the same germ layer. These consistent genes were largely germ layer specific, although mesoderm and endoderm shared more than expected by chance. Ageing profiles were significantly more similar between tissues from the same germ layer than between tissues from different germ layers. Germ layer origin also explained 13.6% of the variation in tissue ageing profiles. Co-expression networks of ageing were organised differently in each germ layer, and 43% of enriched biological processes were specific to one germ layer. However, after grouping related processes, most functional themes were shared by all three, and genes shared between modules of different germ layers were enriched mainly for immune processes. Together, these results indicate that germ layer origin leaves a small but detectable imprint on transcriptomic ageing. However, tissue-specific and shared immune programmes are also prominent.

## 1. Introduction

Ageing is associated with an increased risk of disease and progressive functional decline. However, this decline is not uniform across the body. In humans, physiological function declines at different rates across organ systems (Khan et al., 2017; Tian et al., 2023). Therefore, distinguishing ageing mechanisms that are tissue-specific from those shared across broader groups of tissues could lead to the development of targeted rejuvenation interventions.

The current landscape of ageing biology is dominated by two theories. Damage based theories attribute ageing to the accumulation of molecular and cellular damage. This includes oxidative stress and DNA damage (Harman, 1956; Kirkwood, 1977; Kirkwood, 2008), which drives the progressive deterioration of organs, cells and molecular pathways. This can lead to an increase in disease susceptibility, thereby promoting the onset of age-related pathologies. An alternative theory, proposed by de Magalhães and Church (2005), suggests that pre-established developmental programmes, that were once beneficial for growth and development, become maladaptive later in life, driving an ageing phenotype. This programmatic view implies that ageing is, to some extent, genetically pre-determined, such that the ageing process follows a broadly set trajectory, shared across individuals, but with unique variations due to stochastic effects in each individual. Most likely, ageing reflects a combination of both damage accumulation and programmed processes, and it is within this context that the developmental origin of adult tissues may exert an additional influence on ageing trajectories.

Adult tissues can be classified according to their embryonic germ layer of origin. This can be used to assess similarities in ageing between tissues of shared developmental lineage. During the third week of embryogenesis, gastrulation gives rise to three distinct germ layers, the endoderm, ectoderm and mesoderm (Moore et al., 2013). Cells within these lineages differentiate into distinct tissue types. The ectoderm gives rise to the nervous system and epidermis. The mesoderm produces muscle, connective tissue, blood, the cardiovascular system, kidneys and gonads. The endoderm forms the epithelial lining of the digestive and respiratory tracts and their associated organs, including the liver, pancreas and thyroid (Moore et al., 2013). Germ layer origin is also a major axis of organ transcriptome variation. Across seven organs in seven species, gene expression separated organs by germ layer, with organs most similar early in development and diverging as they mature (Cardoso-Moreira et al., 2019). If tissues from the same lineage display aligned ageing trajectories, this would provide support for the programmed theory of ageing. Such shared ageing pathways across tissues may also represent common targets for rejuvenation therapeutics in anti-ageing research.

Large-scale blood studies have identified many genes that are age-associated, related to immune, metabolic and epigenetic processes (Peters et al., 2015). More recently, multi-tissue analyses using GTEx data have shown that ageing affects gene expression in many tissues, although the size and direction of these effects are mostly tissue-specific (Yamamoto et al., 2022). In contrast, a recent cross species analysis spanning over 25 tissues in four mammals identified transcriptomic hallmarks of ageing conserved across tissues and species including interferon, inflammatory and coagulation programmes (Tyshkovskiy et al., 2026). However, that analysis removed tissue differences to identify universal signals. Intermediate groupings such as germ layer may provide additional information on ageing architecture that sits between tissue-specific and universal ageing.

The Genotype-Tissue Expression (GTEx) project provides bulk RNA-seq data from 54 non-diseased tissue sites. These were collected post-mortem from nearly 1,000 adult donors aged 20–79 years, making it one of the largest resources of healthy human gene expression (GTEx Consortium, 2020). Here, we assigned 24 GTEx tissues to a single germ layer, estimated age effects in each tissue using limma-voom and combined them within each germ layer by random-effects meta-analysis. We then tested whether tissues from the same germ layer age more similarly than tissues from different germ layers. Further analyses explored whether finer developmental relatedness predicts this similarity and whether ageing co-expression networks differ between germ layers. Together, these analyses test whether germ layer origin leaves a detectable signature on transcriptional ageing beyond tissue identity.

## 2. Methods

### Data selection and processing

Bulk RNA-seq count data and matching metadata were obtained from recount3 (Wilks et al., 2021), using GTEx v8 samples (GTEx Consortium, 2020). This provided uniformly pre-processed gene expression data for 32 GTEx tissues. Tissues were grouped according to embryonic germ layer of origin based on standard human embryology (Moore et al., 2013; Supplementary Table S1). These are consistent with germ layer assignments used in previous developmental transcriptome studies (Cardoso-Moreira et al., 2019). Six tissues were excluded from the classification because their bulk tissue composition contains mixed developmental origin or germline content. Two tissues with missing metadata were also removed. The final dataset comprised 24 tissues: mesoderm contained ten tissues with 6,102 samples, endoderm contained ten tissues with 5,232 samples, and ectoderm contained four tissues with 4,069 samples. Full tissue assignments and sample sizes are provided in Supplementary Table S1.

Within each tissue, lowly expressed genes were filtered using filterByExpr() from edgeR. This used a design matrix including age and sex. Genes were retained if they exceeded 0.5 counts per million in a sufficient number of samples for model fitting. Raw counts were then normalised for library size and compositional bias using trimmed mean of M-values. Ages were taken as midpoints from ten-year intervals, providing continuous variables for linear modelling. Non-protein coding genes were also removed to allow for better downstream interpretations.

### Detecting genes with age-related expression changes

Age-associated expression changes were first estimated within each tissue using limma-voom (Law et al., 2014; Ritchie et al., 2015). The voom-transformed expression values for each gene were modelled as a function of age and sex, within each tissue. Age was scaled in units of decades, so the age coefficient represented the estimated log2 fold-change in expression per ten years. Linear modelling was performed using the equation:

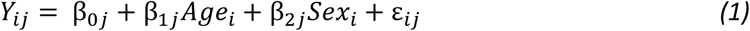

where *Y_ij_* represents the voom-transformed expression of gene *j* in sample *i* for each tissue.

*Age_i_* is the donor age in decades, *Sex_i_* is the donor sex and ε*_ij_* is residual error. Models were fitted with voom precision weights and empirical Bayes moderation of standard errors. Only the age coefficient and its standard error (derived from the moderated t statistic) were carried forward to the meta-analysis. Significance was assessed only after pooling tissues within each germ layer.

### Random-effects meta-analysis within germ layers

Genes with age-effect estimates in at least three tissues within a germ layer were included in the random-effects meta-analysis, performed using the metafor package (Viechtbauer, 2010). This analysis pooled ageing effects for each gene within each germ layer. For tissue t, the random-effects model implemented was:

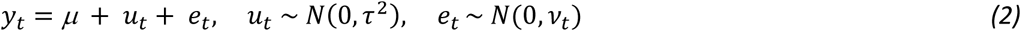

where *y_t_* is the log2 fold change expression per decade, *μ* is the pooled ageing effect (random-effects weighted mean) for that gene within the germ layer, *u_t_* is the tissue-specific deviation from the pooled effect, *τ*^2^ is the between tissue variance, *e_t_* is sampling error and *ν_t_* is sampling variance of the tissue estimate.

Between-tissue variance was calculated using restricted maximum likelihood for most of the genes. For genes with near homogeneous tissue-level effects (defined as a standard deviation of tissue log2 fold-changes below 0.01) the DerSimonian-Laird estimator was used to provide stable variance estimates (DerSimonian and Laird, 1986).

For each gene and germ layer, the meta-analysis provided a pooled ageing effect (μ), standard error, z-statistic, p-value, between-tissue variance (τ²), heterogeneity statistic (I²) and the number of contributing tissues used. Meta-analysis p-values were corrected separately within each germ layer using the Benjamini-Hochberg false discovery rate (FDR) procedure (Benjamini and Hochberg, 1995). Genes with FDR < 0.05 were considered significantly age-associated.

The I² statistic was used to quantify the proportion of total variation attributable to between-tissue variation rather than within-tissue sampling error. Significant genes were stratified by heterogeneity, with low heterogeneity defined as I² ≤ 25% (Higgins et al., 2003). Under a random-effects model, a significant pooled effect can arise when a gene changes strongly with age in only a subset of tissues. Low-heterogeneity genes were therefore used for downstream analyses, as their age effects were consistent across the tissues of a germ layer.

### Low-heterogeneity gene overlap between germ layers

A graph was generated highlighting the gene intersections between the three germ layers. Pairwise enrichment of overlap between germ layers was assessed using two-sided Fisher’s exact tests, based on shared gene identity. To account for direction of effect, the tests were repeated separately for genes upregulated and downregulated with age. Background genes were defined as all genes tested in the germ layer meta-analysis. P-values were adjusted using the Benjamini-Hochberg procedure within each set of tests (Supplementary Tables S3 and S4).

### Ageing and senescence database comparison

The low-heterogeneity genes were compared against curated ageing and senescence resources GenAge and CellAge (Tacutu et al., 2018; Avelar et al., 2020). These were obtained from the Human Ageing Genomic Resource. Ensembl IDs were converted to Entrez IDs using org.Hs.eg.db. Both low-heterogeneity genes and all age-associated genes (FDR < 0.05) were also compared with the 695 genes significantly associated with age in the same direction across the mouse, rat, macaque and human multi-tissue analyses of Tyshkovskiy et al. (2026). Overlap and direction concordance were tested using one-sided Fisher’s exact and binomial tests, respectively, within the genes present in both datasets. Representative genes for Table 1 were selected from the low-heterogeneity genes based on effect size, FDR, number of tissues and database overlap.

**Table 1.** Representative low-heterogeneity ageing genes.

| Germ layer | Gene | log2FC/decade | % change/decade | FDR | I² (%) | n tissues | Overlap |
|---|---|---|---|---|---|---|---|
| ECTODERM | *PF4* | 0.506 | 42.0 | 4.31E-11 | 0.8 | 4 | No GenAge/CellAge overlap |
| ECTODERM | *AHSP* | 0.404 | 32.3 | 5.01E-08 | 0.0 | 4 | No GenAge/CellAge overlap |
| ECTODERM | *KIR2DL3* | 0.368 | 29.1 | 1.57E-06 | 0.0 | 4 | No GenAge/CellAge overlap |
| ECTODERM | *CD40LG* | 0.221 | 16.5 | 8.17E-07 | 0.0 | 4 | CellAge overlap |
| ECTODERM | *GH1* | 0.166 | 12.2 | 4.77E-02 | 0.0 | 4 | GenAge overlap |
| ENDODERM | *OSTN* | 0.291 | 22.4 | 2.85E-10 | 0.4 | 10 | No GenAge/CellAge overlap |
| ENDODERM | *IFIT1B* | 0.270 | 20.6 | 5.01E-05 | 24.8 | 10 | No GenAge/CellAge overlap |
| ENDODERM | *BMP4* | 0.087 | 6.2 | 7.10E-15 | 14.7 | 10 | CellAge overlap |
| ENDODERM | *FGF21* | 0.134 | 9.8 | 6.46E-03 | 0.0 | 6 | GenAge + CellAge overlap |
| MESODERM | *IZUMO1R* | 0.325 | 25.2 | 4.93E-08 | 0.0 | 8 | No GenAge/CellAge overlap |
| MESODERM | *MAS1L* | 0.302 | 23.3 | 1.04E-05 | 17.8 | 9 | No GenAge/CellAge overlap |
| MESODERM | *OR51J1* | 0.260 | 19.8 | 6.92E-11 | 0.0 | 9 | No GenAge/CellAge overlap |
| MESODERM | *TFAP2A* | 0.106 | 7.6 | 3.77E-11 | 2.1 | 10 | GenAge overlap |
| MESODERM | *HMGA2* | 0.106 | 7.6 | 1.02E-03 | 0.0 | 10 | CellAge overlap |

### Tissue ageing profile distance analysis

To test whether tissues from the same germ layer age more similarly, a tissue × gene matrix of age effects was constructed. To avoid selecting genes using the germ layer labels being tested, genes were selected by a random-effects meta-analysis pooling all 24 tissues (same model and estimators as above), retaining 6,534 genes present in at least three tissues with FDR < 0.05. The matrix contained the limma-voom age coefficients for these genes in each tissue and missing values were not imputed. Tissue dissimilarity was defined as one minus the Spearman correlation between age-effect profiles:

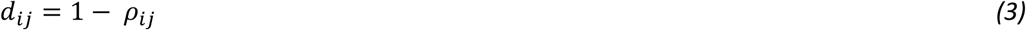

Germ layer structure was tested by PERMANOVA (adonis2, vegan), and differences in dispersion with betadisper (Anderson, 2001; Oksanen et al., 2024). Mean within- and between-germ layer distances were also compared by permuting germ layer labels 9,999 times. The one-sided p-value was the proportion of label arrangements, including the observed one, with a within-minus-between difference as small as or smaller than observed.

To explore whether germ layer also structures baseline expression, and whether this changes with age, we repeated these test on expression profiles from matched young (20–49 years) and old (60–79 years) donors in 21 tissues.

### Developmental-distance analysis

To test whether finer developmental relatedness was associated with tissue ageing similarity, a developmental ontology tree was constructed for the 24 retained tissues. Tissues were further grouped into sub-branches reflecting embryological lineage (neuroectoderm, neural crest and surface ectoderm; paraxial, intermediate and lateral mesoderm; foregut, midgut and hindgut/urogenital sinus endoderm). This was defined using standard human embryology (Moore et al., 2013). Tissues spanning two sub-lineages were placed on their own branches. This included the pituitary, whose anterior and posterior lobes derive from oral ectoderm and neuroectoderm respectively. GTEx colon samples are separated as they span the midgut– hindgut boundary (Fig. 1; Supplementary Table S2). The tree was specified as an ultrametric hierarchy with equal branch lengths at each split, so that distances reflected ordinal developmental relatedness rather than developmental time. Pairwise developmental distances were calculated using cophenetic path length and compared with the tissue ageing-profile distance matrix using a Mantel test with Spearman correlation and 9,999 permutations (Mantel, 1967).

**Figure 1:**
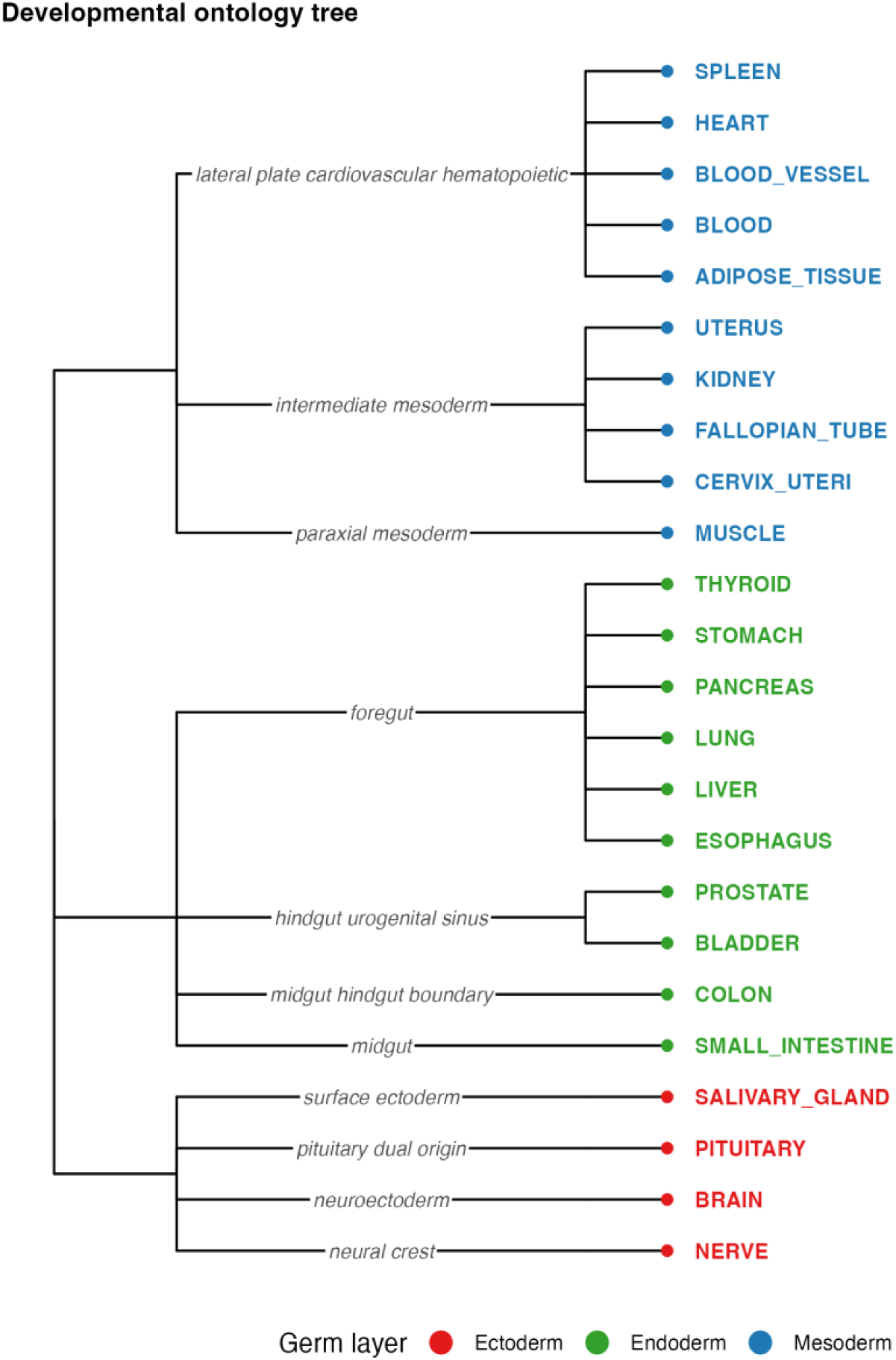
Developmental ontology tree for the 24 tissues. Tips are coloured by germ layer (ectoderm, red; mesoderm, blue; endoderm, green). Sub-branch labels indicate the developmental sub-lineage used to construct the tree. Branch lengths are equal at every split.

### Weighted gene co-expression network analysis

Weighted gene co-expression network analysis (WGCNA) was performed separately within each germ layer (Langfelder and Horvath, 2008). Input matrices were generated from the filtered protein-coding count data used for the initial age-effect analysis. Samples without complete age or sex metadata were removed, as age was later used to test module relationships. For each tissue, counts were normalised using TMM and transformed using limma-voom with a design matrix including sex only. This ensured that age did not contribute to the initial co-expression structure.

Tissue expression matrices were combined within each germ layer using the intersection of genes present across the retained tissues. To remove systematic differences between tissues, residual expression matrices were generated by regressing expression on tissue identity and sex, comparable to the within-tissue centring used in previous multi-tissue ageing network analyses (Tyshkovskiy et al., 2026):

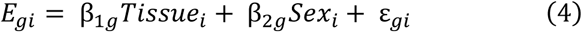

where *E_gi_* is the voom-transformed expression of gene *g* in sample *i*. Tissue*_i_* is the tissue identity of sample *i*, Sex*_i_* is donor sex, and *ε_gi_* is the residual expression value used for WGCNA. Within each germ layer, the top 50% most variable genes by residual variance were retained for network construction. This resulted in 9,456 genes in 4,025 samples for ectoderm, 9,173 genes in 6,066 samples for mesoderm, and 9,280 genes in 5,200 samples for endoderm.

Gene co-expression similarity was calculated using Pearson correlation across samples within each germ layer. Signed weighted networks were constructed using soft-thresholding powers selected from scale-free topology and mean-connectivity diagnostics. The final soft-threshold powers were 8 for ectoderm, 6 for mesoderm and 5 for endoderm. Networks were constructed using signed topological overlap, a minimum module size of 30 genes, dynamic tree cutting, and eigengene-based module merging with a merge cut height of 0.25. No maximum block-size restriction was applied to network construction. Module eigengenes were calculated as the first principal component of each module expression matrix. Grey or unassigned modules were retained during network construction but excluded from downstream enrichment and cross-germ layer overlap analyses.

Module age associations were tested after network construction. For each module, the module eigengene was modelled as a function of age, tissue and sex:

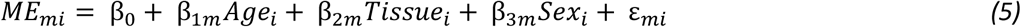

where *ME_mi_* is the eigengene value of module m in sample i, *Age_i_* is donor age in decades, *Tissue_i_* is tissue identity, *Sex_i_* is donor sex and ε*_mi_* is residual error. The age coefficient β_1*m*_ therefore represents the estimated change in module eigengene activity per decade. P-values for the age coefficient were adjusted for multiple testing within each germ layer using the Benjamini–Hochberg false discovery rate procedure. Modules with FDR < 0.05 were considered age-associated. Because the sign of a principal component is arbitrary, module interpretation was based on the presence and magnitude of age association rather than eigengene direction.

Module architecture was summarised using the total number of modules (excluding the grey module), the number of age-associated modules and the distribution of module sizes. Eigengene correlation networks were constructed from age-associated modules by calculating pairwise Pearson correlations between module eigengenes within each germ layer. Edges were retained where |r| ≥ 0.5, with edge sign indicating positive or negative eigengene correlation. Nodes represented age-associated modules and node size represented module size.

Functional enrichment was performed for non-grey age-associated modules using GO Biological Process and Reactome pathway annotations (Ashburner et al., 2000; Yu et al., 2012; Yu and He, 2016; Gillespie et al., 2022). The enrichment background was defined separately for each germ layer as all genes assigned to WGCNA modules in that network. Enrichment analyses were performed using compareCluster() from clusterProfiler, with GO Biological Process tested using enrichGO() and Reactome pathways tested using enrichPathway() from ReactomePA. Benjamini-Hochberg correction was applied, and terms with adjusted p < 0.05 were retained. Since many enriched GO terms are redundant, significant GO Biological Process terms from the age-associated modules of all three germ layers were grouped by semantic similarity using rrvgo (Sayols, 2023), with the Rel similarity measure and the default threshold of 0.7. The most significant term in each group was used as the theme name, and a theme was counted as present in a germ layer if any of its terms was enriched there.

Overlap between age-associated modules from different germ layers (excluding grey modules) was tested using one-sided Fisher’s exact tests on gene membership, with Benjamini–Hochberg correction. The background was the 7,398 genes present in all three germ layer networks. Genes from significantly overlapping pairs (FDR < 0.05) were tested for GO Biological Process and Reactome enrichment against the same background.

## 3. Results

### Random-effects meta-analysis identifies distinct ageing genes across germ layers

Significant age-associated genes were detected in all three germ layers. Before heterogeneity filtering, endoderm had the most (4,805), followed by mesoderm (4,520) and ectoderm (1,744) (Fig. 2A). This order reversed after restricting to genes with low between-tissue heterogeneity (I² ≤ 25%), whose age effects were consistent across tissues of a germ layer. Ectoderm retained 1,035 genes (59.4% of its significant genes), mesoderm 456 (10.1%) and endoderm 409 (8.5%) (Fig. 2B). Most low-heterogeneity genes were upregulated with age in mesoderm (70.6%) and endoderm (66.0%), whereas ectoderm was more evenly split (54.7% up) (Fig. 2C). Most genes had modest effects per decade, although each germ layer included a subset with larger effects (Fig. 2D).

**Figure 2:**
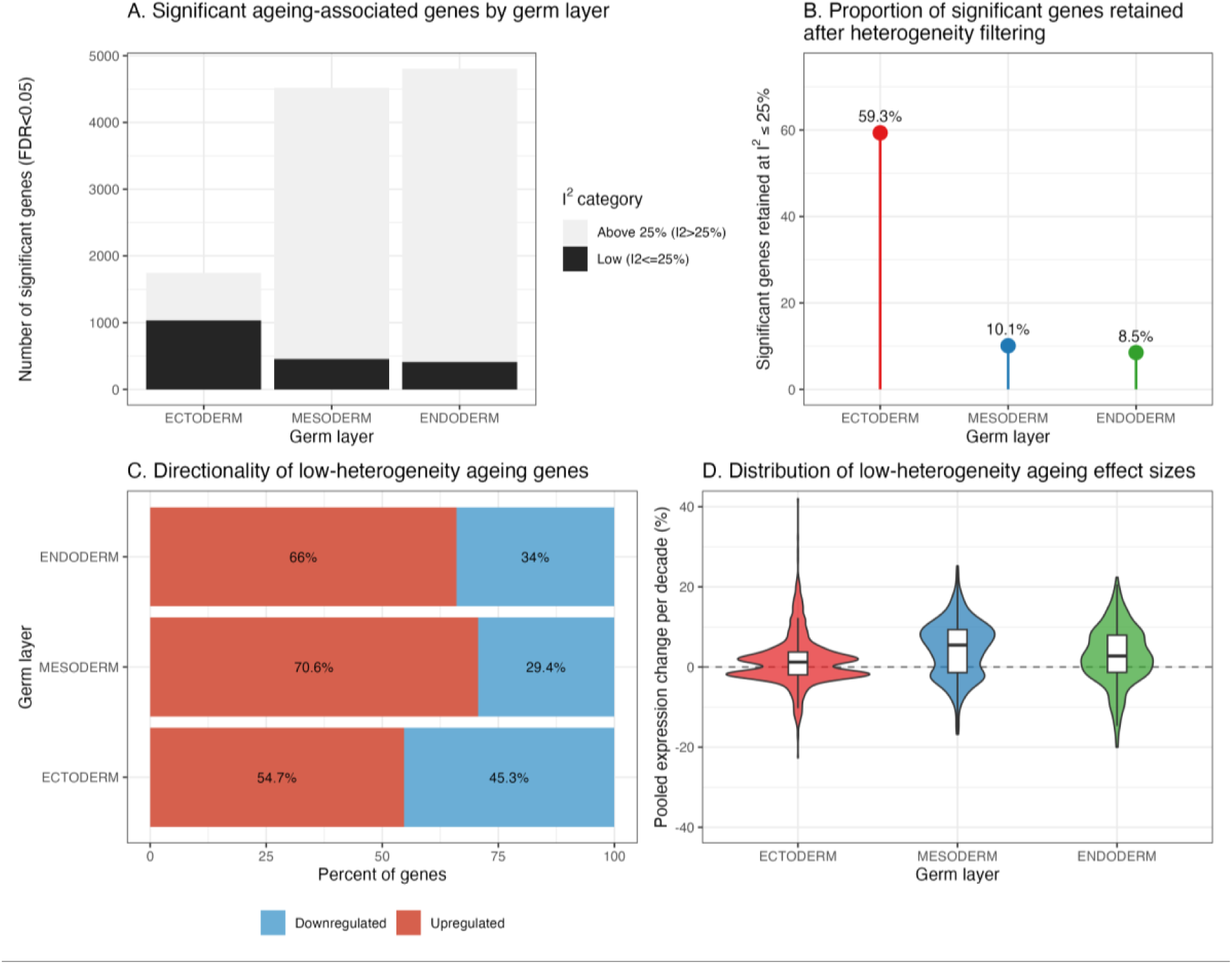
Germ layer meta-analysis identifies age-associated genes. A. Total number of significant age-associated genes (FDR<0.05) per germ layer, split into low-heterogeneity (I²≤25%) and higher-heterogeneity (I²>25%) genes. B. Proportion of significant genes retained after low-heterogeneity filtering. C. Directionality of low-heterogeneity genes. D. Distribution of expression change per decade among low-heterogeneity genes.

### Low-heterogeneity ageing genes are largely lineage-specific and mostly uncatalogued

Most low-heterogeneity genes were specific to one germ layer (989 ectoderm, 403 mesoderm, 360 endoderm). Mesoderm and endoderm shared 27 genes, ectoderm and mesoderm 24, and ectoderm and endoderm 20. Two genes, *CACNG7* and *KLKB1*, were shared by all three (Fig. 3). Only the mesoderm–endoderm overlap was greater than expected by chance (Fisher’s exact test, FDR = 4.9 × 10⁻⁷; Supplementary Table S3). When genes were split by direction of change, overlap was significant for genes upregulated in both mesoderm and endoderm (FDR = 1.3 × 10⁻¹¹). This was also true for genes downregulated in both mesoderm and ectoderm (FDR = 1.3 × 10⁻³; Supplementary Table S4). *KLKB1* was upregulated with age in all three germ layers, whereas *CACNG7* was downregulated in ectoderm but upregulated in mesoderm and endoderm.

**Figure 3:**
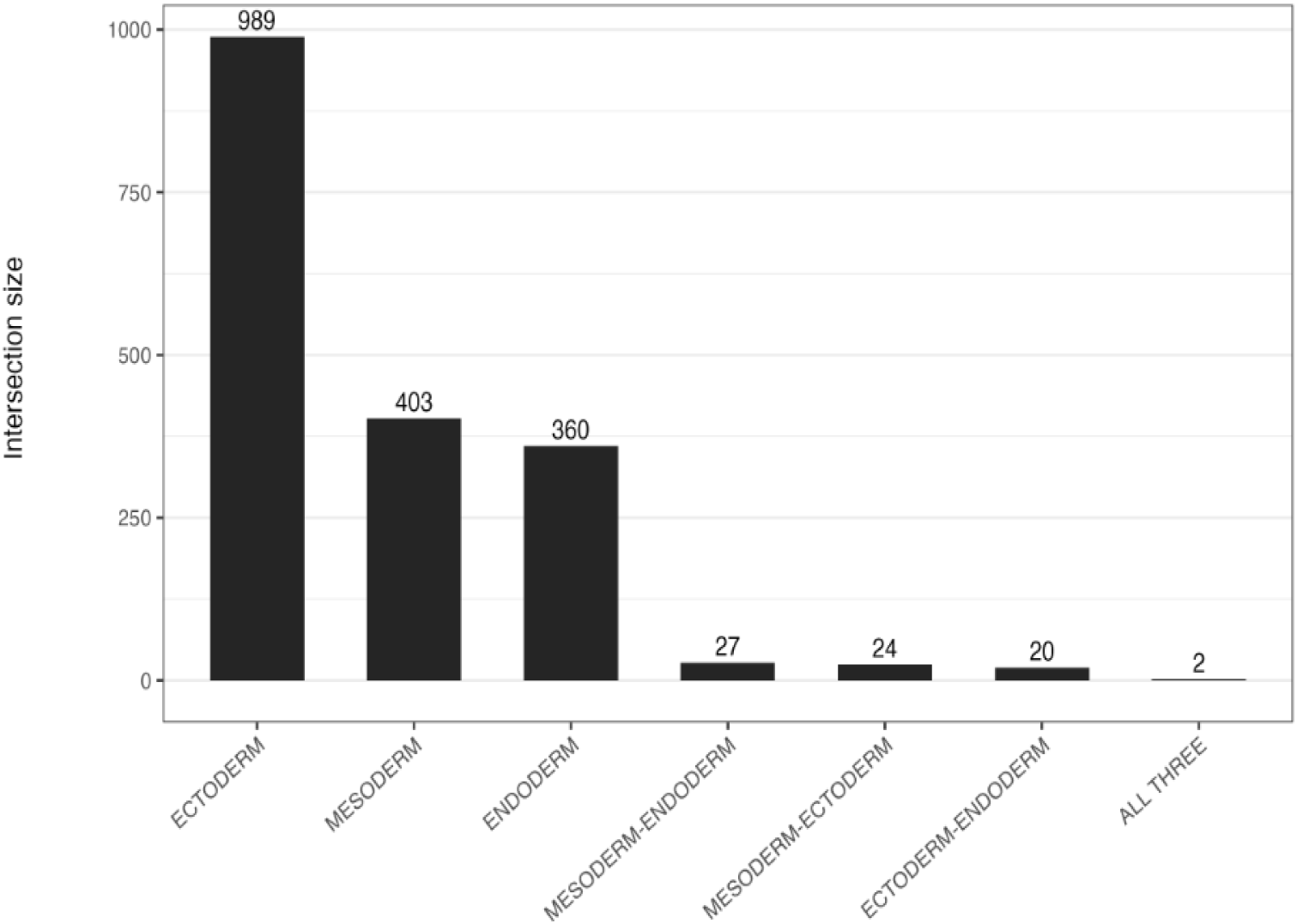
Bar chart showing intersections between significant low-heterogeneity ageing genes identified in ectoderm, mesoderm and endoderm. The filled points indicate the germ layers represented in each set.

Only 79 low-heterogeneity genes (4.3%) overlapped GenAge or CellAge, with 62 in ectoderm, 11 in endoderm and 7 in mesoderm. Ectodermal genes overlapped at the expected rate (odds ratio 1.10, FDR = 0.70). Mesodermal and endodermal genes were under represented (odds ratio 0.26, FDR = 7.3 × 10⁻⁵ and odds ratio 0.46, FDR = 0.025, respectively; Supplementary Table S5). This indicates that most consistent germ layer ageing genes have not previously been catalogued as ageing or senescence genes. Overlapping genes included *CD40LG* and *GH1* in ectoderm, *FGF21* and *BMP4* in endoderm, and *TFAP2A* and *HMGA2* in mesoderm. Neither low-heterogeneity genes nor all age-associated genes overlapped the 695 genes that change consistently with age across mouse, rat, macaque and human tissues (Tyshkovskiy et al., 2026) more than expected by chance. However, among all age-associated genes, the direction of change largely agreed where genes overlapped, particularly in mesoderm (70.8% concordant, binomial p = 2.8 × 10⁻⁸; Supplementary Table S5). Several of the strongest ectodermal genes are markers of circulating cells, including *PF4* (platelets and megakaryocytes), *AHSP* (erythroid cells) and *KIR2DL3* (natural killer cells). Other representative genes with large effects are shown in Table 1.

### Tissue-tissue similarity is partly structured by germ layer

Tissue ageing-profile distances were analysed to determine whether broader ageing patterns retained developmental lineage structure. Distances were calculated from 6,534 significantly age associated genes in a meta-analysis pooling all 24 tissues. This was visualized by hierarchical clustering (Supplementary Fig. 1) and principal coordinates analysis (PCoA; Fig. 4). The PCoA showed partial germ layer organisation. The ectoderm centroid was more separated from the others, whereas mesodermal and endodermal tissues were substantially intermixed (Fig. 4).

**Figure 4:**
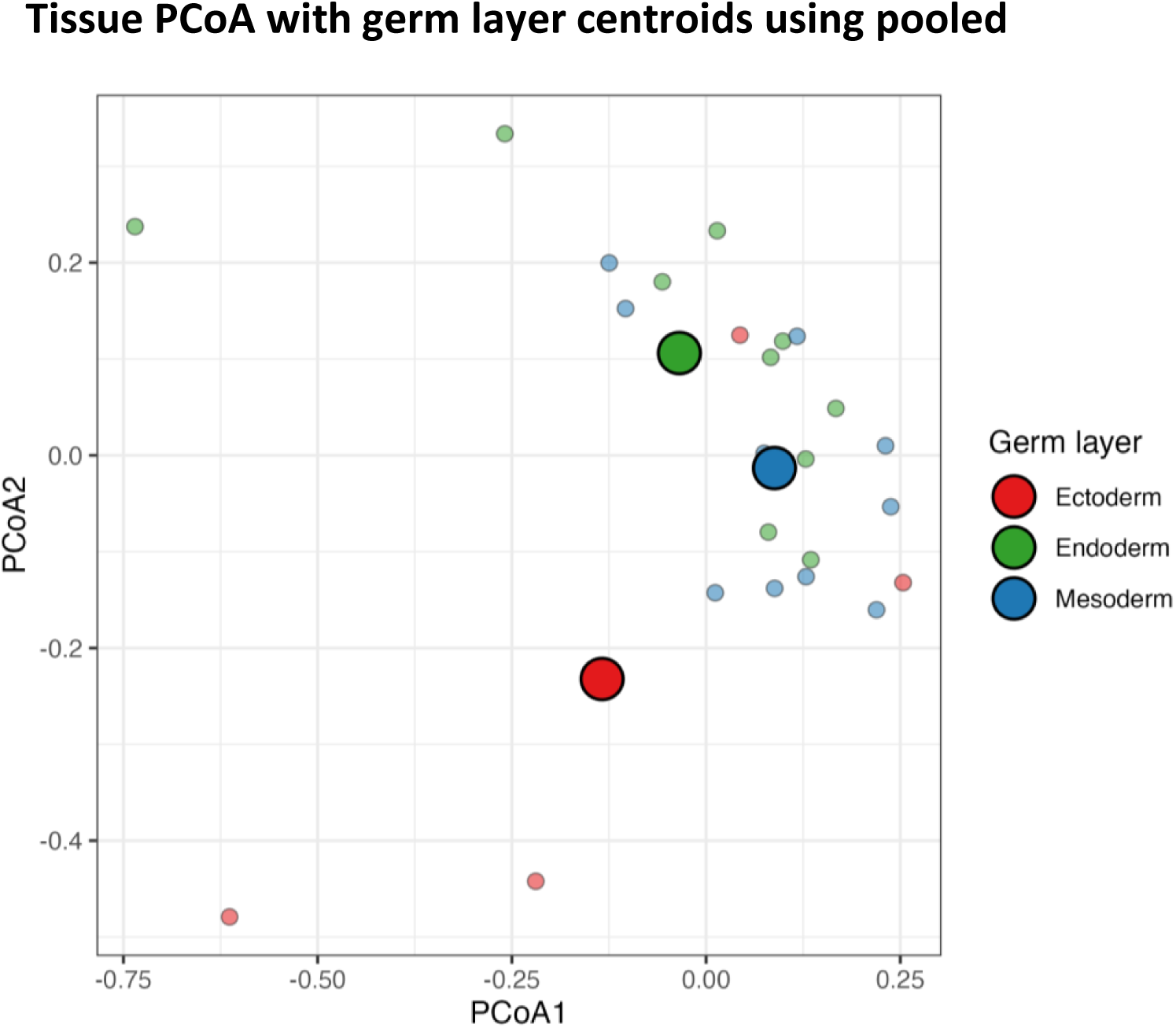
Principal coordinates analysis (PCoA) of tissue ageing-profile distances (1 − Spearman ρ) for the 24 tissues, based on 6,534 genes significantly age-associated (FDR < 0.05) in a meta-analysis pooling all 24 tissues. Small, outlined points show individual tissues and large points show germ layer centroids. Both are coloured by germ layer.

PERMANOVA supported a small germ layer effect, with germ layer explaining 13.6% of the variation in tissue ageing-profile distances (p = 0.035). No significant difference in dispersion was detected among germ layers (betadisper p = 0.355), indicating that this result was not driven by unequal spread around germ layer centroids. Tissues from the same germ layer also had lower ageing-profile distances than tissues from different germ layers (within − between difference = −0.057, p = 0.029). A Mantel test comparing developmental distances from the ontology tree with ageing-profile distances showed a positive correlation (Spearman r = 0.155, p = 0.019). This suggests that finer embryological relatedness explains a small but detectable component of tissue ageing variation.

Germ layer also structured baseline expression in both young (PERMANOVA R² = 0.184, p = 0.011) and old donors (R² = 0.179, p = 0.013). This did not differ between age groups in the primary analysis (p = 0.29), and no tissue moved significantly closer to or further from its own germ layer with age (Supplementary Table S6).

### Co-expression module architecture differs between germ layers

Co-expression networks differed between germ layers in the number of modules and the proportion associated with age (Fig. 5A). A module was considered age-associated when its eigengene changed significantly with age after adjusting for tissue and sex (FDR < 0.05), meaning that many of its genes change together with age. Mesoderm had the most modules (23, excluding the unassigned grey module), 17 of them age-associated, compared with 16 (11 age-associated) in ectoderm and 12 (10 age-associated) in endoderm. Endoderm therefore had the highest proportion of age-associated modules (83.3%), followed by mesoderm (73.9%) and ectoderm (68.8%) (Fig. 5A,B).

**Figure 5:**
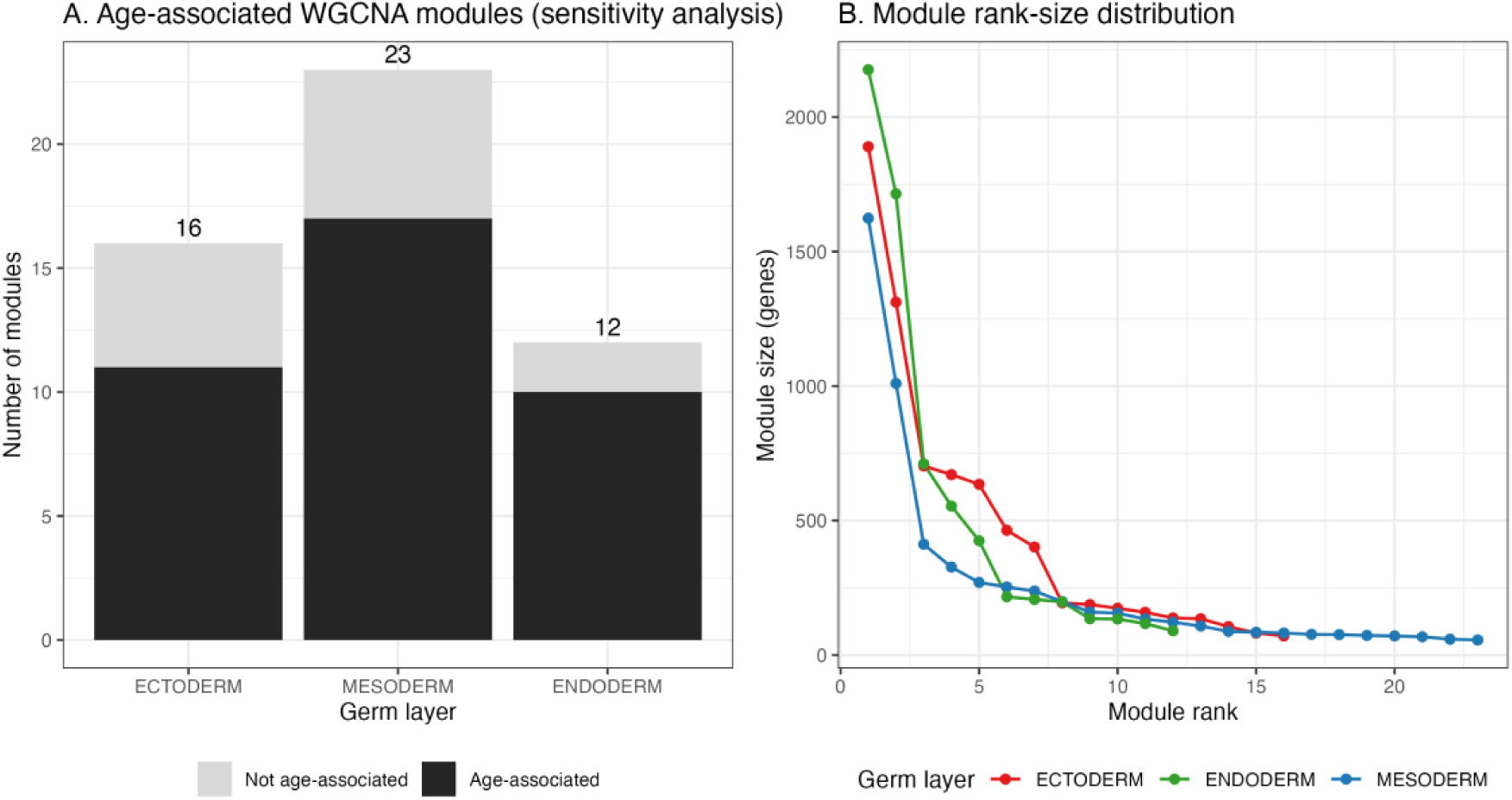
WGCNA module architecture across germ layers. (A) Number of co-expression modules per germ layer. (B) Module rank-size distribution. Module size (number of genes) plotted against rank for each germ layer (ectoderm, red; mesoderm, blue; endoderm, green).

The three germ layers also differed in how their age-associated modules were connected to one another (Fig. 6). In ectoderm, all 11 age-associated modules formed a single connected network at |r| ≥ 0.5. This suggests that ectodermal ageing behaves as one broadly coordinated transcriptional programme rather than several independent processes. Endoderm was similarly cohesive, with nine of its ten age-associated modules forming a connected structure and only one module acting independently. Mesoderm showed the opposite pattern. Although it contained the most age-associated modules, these fragmented into four separate clusters, with three modules entirely uncorrelated with any other age-associated module.

**Figure 6:**
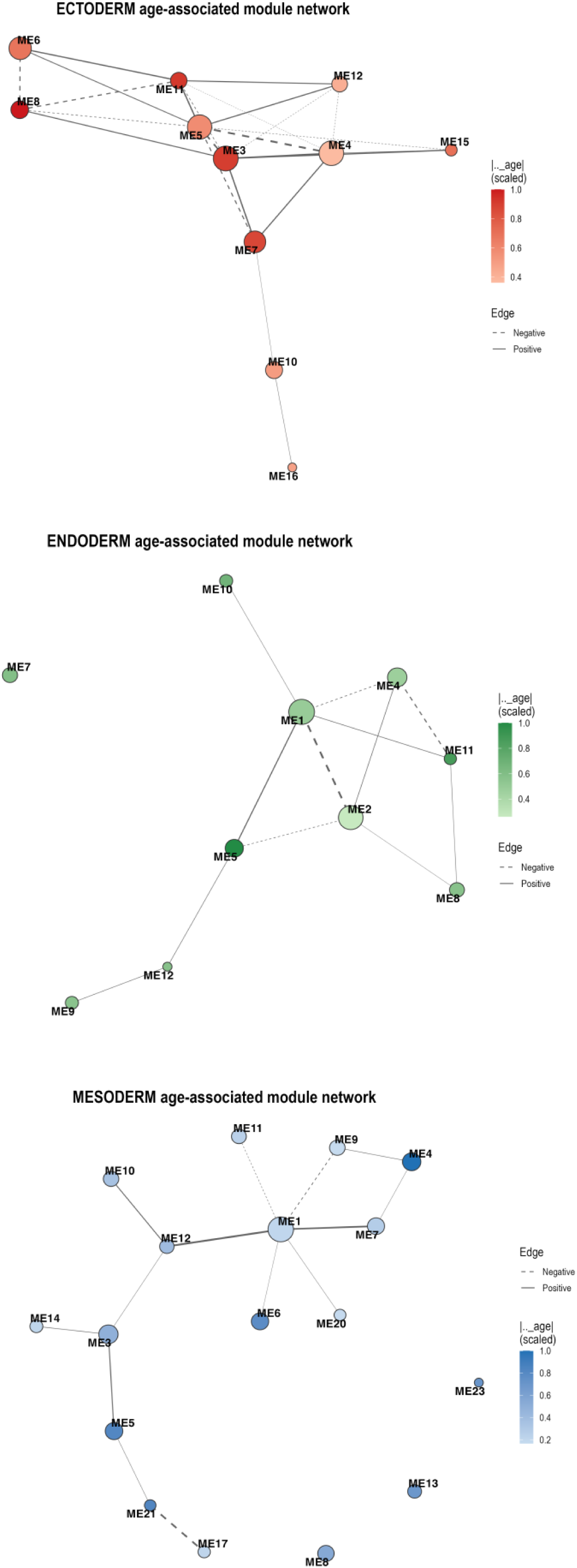
Eigengene correlation networks of age-associated WGCNA modules within each germ layer. Each node is an age-associated module. Node size represents module size and fill intensity represents the magnitude of the module’s age coefficient (darker modules showing a stronger age effect). Edges connect module pairs with eigengene correlation |r| ≥ 0.5. Solid lines show positive and dashed lines negative correlations.

### Functional enrichment among age-associated modules

Within each germ layer, most enriched GO Biological Process terms were found in only one module (ectoderm 78%, endoderm 86%, mesoderm 68%). This suggests that age-associated modules capture largely distinct processes. Across germ layers, 43% of all terms (1,159 of 2,681) were enriched in only one germ layer, but many were closely related to terms enriched in another. When related terms were grouped into 172 themes by semantic similarity, more than half (54%) were shared by all three germ layers and only 17% were specific to one (Fig. 7).

**Figure 7:**
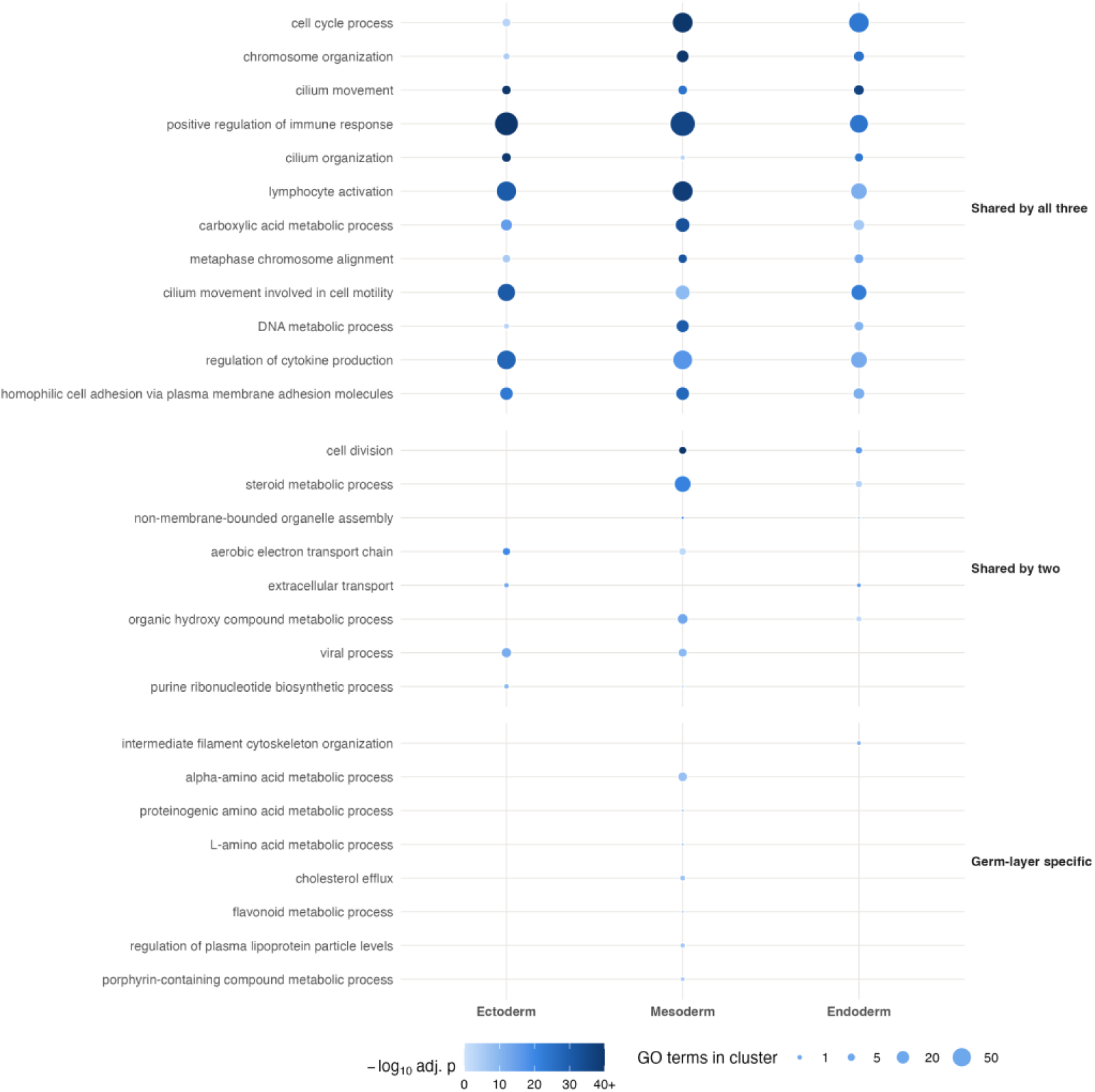
Semantic reduction of GO Biological Process terms enriched in age-associated modules. Enriched terms (adjusted p < 0.05) were grouped by semantic similarity (rrvgo; Rel measure, threshold 0.7). Rows show representative themes enriched in all three, two or one germ layer. Colour intensity shows the strongest enrichment (−log₁₀ adjusted p, capped at 40); size shows the number of enriched GO terms in the theme within each germ layer.

Shared themes included immune regulation, cell cycle and cilium organisation. Germ layer specific themes were mainly metabolic in mesoderm (amino acid metabolism, cholesterol efflux) and structural in endoderm (intermediate filament organisation).

Although co-expression structure differed between germ layers, the age-associated modules were not composed of entirely separate genes. Of the 467 pairs of age-associated modules from different germ layers, 341 (73.0%) shared at least one gene and 49 (10.5%) overlapped more than expected by chance. The genes in these significantly overlapping pairs (n = 2,167) were mainly involved in immune processes. The top GO Biological Process term was regulation of immune response (adjusted p = 3.3 × 10⁻¹³), and the four top Reactome pathways were all immune-related. Vascular development and cell-cycle processes were also enriched.

## 4. Discussion

This study tested whether embryonic germ layer origin influences adult transcriptomic ageing. Across the germ layer meta-analysis, tissue ageing profile distances and co-expression network analysis, germ layer explained a modest but measurable proportion of ageing variation (Figs. 2-7). These results are inconsistent with a model in which transcriptomic ageing is either entirely tissue-specific or fully shared across tissues. They suggest that developmental origin imposes a measurable constraint on ageing, on top of which tissue-specific effects predominate.

This pattern was clearest in the germ layer meta-analysis (Fig. 2). Endoderm and mesoderm had the most significant age-associated genes but retained relatively few with consistent effects across tissues, which may reflect the functional and cellular diversity of these groups. Ectoderm showed the opposite pattern, although it includes only four tissues. These results fit with reports that most age effects in GTEx are tissue-specific (Yamamoto et al., 2022) and that ageing involves both shared and tissue-specific changes across mammalian tissues (Palmer et al., 2021). More recently, a conserved core of ageing signatures shared across tissues and species was identified (Tyshkovskiy et al., 2026). Germ layer origin may account for part of the variation between these universal and tissue-specific effects. Overlap of consistent genes between germ layers was limited, but mesoderm and endoderm shared more genes than expected by chance. These were mostly upregulated with age, suggesting that these lineages share part of their ageing programme. Most of these genes were not previously catalogued in GenAge or CellAge and did not preferentially overlap conserved cross-species ageing genes. However, across all age-associated genes, the direction of change largely agreed with the conserved genes where they overlapped.

Tissue ageing-profile distances showed a corresponding pattern (Fig. 4). Tissues from the same germ layer had more similar ageing profiles than tissues from different germ layers, even when genes were selected without reference to germ layer. This is the most direct evidence for a germ layer effect in our data, but the effect was modest (R² = 0.136). The PCoA showed substantial overlap between mesodermal and endodermal tissues, and the association between developmental and ageing-profile distances was weak (Mantel r = 0.155). A modest imprint is expected given how organ transcriptomes develop. Germ layer origin is the main axis of organ transcriptome variation early in development, but organ transcriptomes diverge progressively as tissues mature (Cardoso-Moreira et al., 2019). Furthermore, partial loss of tissue identity and convergence between tissues have been reported during ageing (Yamamoto et al., 2022). A residual germ layer structure in adult ageing profiles is therefore consistent with lineage effects that are strongest early in development and increasingly overlaid by tissue specific programmes. Germ layer structure in baseline expression was also similar in young and old donors, suggesting that this imprint is stable through adulthood rather than changing with age.

Network analysis supported the same conclusion. Module architecture differed between germ layers (Fig. 5), but module number depends on network parameters, so these are best interpreted as relative differences between germ layers analysed in the same way rather than absolute properties. Functionally, the germ layers largely converged. Although 43% of individual GO terms were enriched in only one germ layer, after grouping related terms more than half of the themes (54%) were shared by all three. These included immune regulation, cell cycle and cilium organisation, whereas only 17% of themes were specific to one germ layer (Fig. 7). Fewer germ layer specific themes were present. These were mainly metabolic in mesoderm and structural in endoderm. Genes shared between significantly overlapping module pairs were also strongly enriched for immune processes. Adult tissues therefore appear to age through a largely shared, immune-related transcriptional programme, with a smaller set of lineage-specific pathways.

These findings should be considered against the two theories of ageing outlined in the Introduction. Under a programmed view, tissues sharing a developmental origin might be expected to share ageing trajectories (de Magalhães and Church, 2005). Our analyses offer support for this, as germ layer origin leaves a detectable imprint on adult ageing profiles. However, this signal is small relative to tissue-specific effects, and the dominant shared theme is immune and inflammatory. Notably, in mice, transcriptomic age declines during early embryogenesis before rising from around mid-organogenesis. This decline occurs in lineages derived from all three germ layers (Tyshkovskiy et al., 2026). Ageing trajectories may therefore begin close to the point at which germ layer lineages are specified, allowing lineage-specific programmes to shape later ageing. Overall, our results fit a model in which ageing reflects both damage accumulation and developmental programmes and germ layer origin contributes a modest component.

This study has several limitations. First, bulk RNA-seq measures a mixture of cell types whose proportions may change with age. Several of the strongest ectodermal genes are blood-cell markers (*PF4, AHSP, KIR2DL3*), so their increase may reflect more blood or immune cells in older tissues rather than changes in neural cells. Single-cell or deconvolution analyses would help separate lineage effects from changes in cell composition. Second, assigning each tissue to a single germ layer is a simplification of the developmental biology. Most organs contain cells from more than one germ layer. For example, the gut epithelium derives from endoderm, whereas its muscle and connective tissue derive from mesoderm (Moore et al., 2013). Blood, immune and vascular cells, which are mesodermal, are also present in every tissue. This may dilute germ layer specific signals and contribute to the immune and vascular themes shared across germ layers. Third, GTEx is cross-sectional, so it captures differences between age groups rather than changes within individuals over time. Finally, some tissues had modest sample sizes, and the ectoderm meta-analysis included only four tissues.

Overall, embryonic germ layer origin does not appear to be the main driver of adult transcriptomic ageing in humans. However, it leaves a small but consistent developmental signature. This is detectable at the gene, tissue and network level, although tissue-specific ageing programmes predominate.

## Data and code availability

GTEx v8 RNA-seq data are publicly available and were accessed through recount3 (Wilks et al., 2021). All analysis code will be made publicly available on GitHub upon publication.

## Acknowledgements

We sincerely thank the members of the Genomics of Ageing and Rejuvenation Lab for their support, valuable discussions and input throughout this project. The Genotype-Tissue Expression (GTEx) Project was supported by the Common Fund of the Office of the Director of the National Institutes of Health, and by NCI, NHGRI, NHLBI, NIDA, NIMH and NINDS. During the preparation of this work, the authors used Claude (Anthropic) to assist with editing text and code. After using this tool, the authors reviewed and edited the content as needed and take full responsibility for the content of the publication.

## Funding

This work was supported by the Biotechnology and Biological Sciences Research Council (BBSRC) and University of Birmingham funded Midlands Integrative Biosciences Training Partnership (MIBTP) [grant number BB/T00746X/1]. Work in our lab is further supported by grants from LongeCity and Longevity Impetus Grants/Hevolution Foundation.

## Conflict of Interest

JPM is CSO of YouthBio Therapeutics, an advisor/consultant for ElixIRA Pharma, AND Biotech AG, and NOVOS. JPM is also the founder of Magellan Science Ltd, a company providing consulting services in longevity science. TE declares no conflicts of interest.

## References

Anderson, M. J. (2001). A new method for non-parametric multivariate analysis of variance. Austral Ecology, 26(1), 32–46. doi: 10.1111/j.1442-9993.2001.01070.pp.x.

Ashburner, M., Ball, C. A., Blake, J. A., Botstein, D., Butler, H., Cherry, J. M., Davis, A. P., Dolinski, K., Dwight, S. S., Eppig, J. T., Harris, M. A., Hill, D. P., Issel-Tarver, L., Kasarskis, A., Lewis, S., Matese, J. C., Richardson, J. E., Ringwald, M., Rubin, G. M. and Sherlock, G. (2000). Gene ontology: tool for the unification of biology. Nature Genetics, 25(1), 25–29. doi: 10.1038/75556.

Avelar, R. A., Ortega, J. G., Tacutu, R., Tyler, E. J., Bennett, D., Binetti, P., Budovsky, A., Chatsirisupachai, K., Johnson, E., Murray, A., Tejada-Martinez, D., Thornton, D., Fraifeld, V. E., Bishop, C. L. and de Magalhães, J. P. (2020). A multidimensional systems biology analysis of cellular senescence in aging and disease. Genome Biology, 21, 91. doi: 10.1186/s13059-020-01990-9.

Benjamini, Y. and Hochberg, Y. (1995). Controlling the false discovery rate: a practical and powerful approach to multiple testing. Journal of the Royal Statistical Society: Series B, 57(1), 289–300. doi: 10.1111/j.2517-6161.1995.tb02031.x.

Cardoso-Moreira, M., et al. and Kaessmann, H. (2019). Gene expression across mammalian organ development. Nature, 571, 505–509. doi: 10.1038/s41586-019-1338-5.

de Magalhães, J. P. and Church, G. M. (2005). Genomes optimize reproduction: aging as a consequence of the developmental program. Physiology, 20, 252–259. doi: 10.1152/physiol.00010.2005.

DerSimonian, R. and Laird, N. (1986). Meta-analysis in clinical trials. Controlled Clinical Trials, 7(3), 177–188. doi: 10.1016/0197-2456(86)90046-2.

Gillespie, M., Jassal, B., Stephan, R., Milacic, M., Rothfels, K., Senff-Ribeiro, A., Griss, J., Sevilla, C., Matthews, L., Gong, C., Deng, C., Varusai, T., Ragueneau, E., Haider, Y., May, B., Shamovsky, V., Weiser, J., Brunson, T., Sanati, N., Beckman, L., Shao, X., Fabregat, A., Sidiropoulos, K., Murillo, J., Viteri, G., Cook, J., Shorser, S., Bader, G., Demir, E., Sander, C., Haw, R., Wu, G., Stein, L., Hermjakob, H. and D’Eustachio, P. (2022). The Reactome pathway knowledgebase 2022. Nucleic Acids Research, 50(D1), D687–D692. doi: 10.1093/nar/gkab1028.

GTEx Consortium. (2020). The GTEx Consortium atlas of genetic regulatory effects across human tissues. Science, 369(6509), 1318–1330. doi: 10.1126/science.aaz1776.

Harman, D. (1956). Aging: a theory based on free radical and radiation chemistry. Journal of Gerontology, 11(3), 298–300. doi: 10.1093/geronj/11.3.298.

Higgins, J. P. T., Thompson, S. G., Deeks, J. J. and Altman, D. G. (2003). Measuring inconsistency in meta-analyses. BMJ, 327(7414), 557–560. doi: 10.1136/bmj.327.7414.557.

Khan, S. S., Singer, B. D. and Vaughan, D. E. (2017). Molecular and physiological manifestations and measurement of aging in humans. Aging Cell, 16(4), 624–633. doi: 10.1111/acel.12601.

Kirkwood, T. B. L. (1977). Evolution of ageing. Nature, 270, 301–304. doi: 10.1038/270301a0.

Kirkwood, T. B. L. (2008). Understanding ageing from an evolutionary perspective. Journal of Internal Medicine, 263(2), 117–127. doi: 10.1111/j.1365-2796.2007.01901.x.

Langfelder, P. and Horvath, S. (2008). WGCNA: an R package for weighted correlation network analysis. BMC Bioinformatics, 9, 559. doi: 10.1186/1471-2105-9-559.

Law, C. W., Chen, Y., Shi, W. and Smyth, G. K. (2014). voom: precision weights unlock linear model analysis tools for RNA-seq read counts. Genome Biology, 15, R29. doi: 10.1186/gb-2014-15-2-r29.

Mantel, N. (1967). The detection of disease clustering and a generalized regression approach. Cancer Research, 27(2), 209–220.

Moore, K. L., Persaud, T. V. N. and Torchia, M. G. (2013). The Developing Human: Clinically Oriented Embryology. 9th edn. Philadelphia: Saunders/Elsevier.

Oksanen, J., Simpson, G. L., Blanchet, F. G., Kindt, R., Legendre, P., Minchin, P. R., O’Hara, R. B., Solymos, P., Stevens, M. H. H., Szoecs, E., Wagner, H., Barbour, M., Bedward, M., Bolker, B., Borcard, D., Carvalho, G., Chirico, M., De Caceres, M., Durand, S., Evangelista, H. B. A., FitzJohn, R., Friendly, M., Furneaux, B., Hannigan, G., Hill, M. O., Lahti, L., McGlinn, D., Ouellette, M.-H., Ribeiro Cunha, E., Smith, T., Stier, A., Ter Braak, C. J. F. and Weedon, J. (2024). vegan: Community Ecology Package. R package version 2.6–6.1.

Palmer D, Fabris F, Doherty A, Freitas AA, de Magalhães JP. Ageing transcriptome meta-analysis reveals similarities and differences between key mammalian tissues. Aging (Albany NY). 2021 Feb 11;13(3):3313–3341. doi: 10.18632/aging.202648. Epub 2021 Feb 11. PMID: 33611312; PMCID: PMC7906136.

Peters, M. J., Joehanes, R., Pilling, L. C., Schurmann, C., Conneely, K. N., Powell, J., Reinmaa, E., Sutphin, G. L., Zhernakova, A., Schramm, K., Wilson, Y. A., Kobes, S., Tukiainen, T., NABEC/UKBEC Consortium, Ramos, Y. F., Göring, H. H., Fornage, M., Liu, Y., Gharib, S. A., Stranger, B. E., De Jager, P. L., Aviv, A., Levy, D., Murabito, J. M., Munson, P. J., Huan, T., Hofman, A., Uitterlinden, A. G., Rivadeneira, F., van Rooij, J., Stolk, L., Broer, L., Verbiest, M. M. P. J., Jhamai, M., Arndt, P. F., Wang, L., Willemsen, G., Hottenga, J. J., Jansen, R., Penninx, B. W., Emilsson, V., Hansen, K. D., Kohanski, R. A., Ronningen, K., Harris, J. R., Ilic, N., Cusack, W., Landefeld, C. S., Lam, M. P. Y., Chan, C., Nöthen, M. M., Nikolaus, S., Franke, A., Balliu, B., Vermeulen, S. H., Kiemeney, L. A., van der Harst, P., Ikram, M. A., Boomsma, D. I., Pers, T. H., Powell, J. E., Montgomery, G. W., Visscher, P. M., Wray, N. R., Hemani, G., Franke, L., van Meurs, J. B. J., van den Berg, L. H., Ferrucci, L. and Slagboom, P. E. (2015). The transcriptional landscape of age in human peripheral blood. Nature Communications, 6, 8570. doi: 10.1038/ncomms9570.

Ritchie, M. E., Phipson, B., Wu, D., Hu, Y., Law, C. W., Shi, W. and Smyth, G. K. (2015). limma powers differential expression analyses for RNA-sequencing and microarray studies. Nucleic Acids Research, 43(7), e47. doi: 10.1093/nar/gkv007.

Sayols, S. (2023). rrvgo: a Bioconductor package for interpreting lists of Gene Ontology terms. microPublication Biology. doi: 10.17912/micropub.biology.000811.

Tacutu, R., Thornton, D., Johnson, E., Budovsky, A., Barardo, D., Craig, T., Diana, E., Lehmann, G., Toren, D., Wang, J., Fraifeld, V. E. and de Magalhães, J. P. (2018). Human Ageing Genomic Resources: new and updated databases. Nucleic Acids Research, 46(D1), D1083–D1090. doi: 10.1093/nar/gkx1042.

Tian, Y. E., et al. and Zalesky, A. (2023). Heterogeneous aging across multiple organ systems and prediction of chronic disease and mortality. Nature Medicine, 29, 1221–1231. doi: 10.1038/s41591-023-02296-6.

Tyshkovskiy, A., Kholdina, D., Davitadze, M., Molière, A., et al. and Gladyshev, V. N. (2026). Universal transcriptomic hallmarks of mammalian ageing and mortality. Nature, 654, 173–188. doi: 10.1038/s41586-026-10542-3.

Viechtbauer, W. (2010). Conducting meta-analyses in R with the metafor package. Journal of Statistical Software, 36(3), 1–48. doi: 10.18637/jss.v036.i03.

Wilks, C. et al. (2021). recount3: summaries and queries for large-scale RNA-seq expression and splicing. Genome Biology, 22, 323. doi: 10.1186/s13059-021-02533-6.

Yamamoto, R., Chung, R., Vazquez, J. M., Sheng, X., Steinberg, J. D., Ioannidis, N. M., Sudmant, P. H. and Lappalainen, T. (2022). Tissue-specific impacts of aging and genetics on gene expression patterns in humans. Nature Communications, 13, 5803. doi: 10.1038/s41467-022-33509-0.

Yu, G. and He, Q. Y. (2016). ReactomePA: an R/Bioconductor package for Reactome pathway analysis and visualization. Molecular BioSystems, 12(2), 477–479. doi: 10.1039/C5MB00663E.

Yu, G., Wang, L. G., Han, Y. and He, Q. Y. (2012). clusterProfiler: an R package for comparing biological themes among gene clusters. OMICS, 16(5), 284–287. doi: 10.1089/omi.2011.0118.

