## Supplementary Figure 1 for "The Impact of Germ Layer Origin on the Ageing Transcriptome"

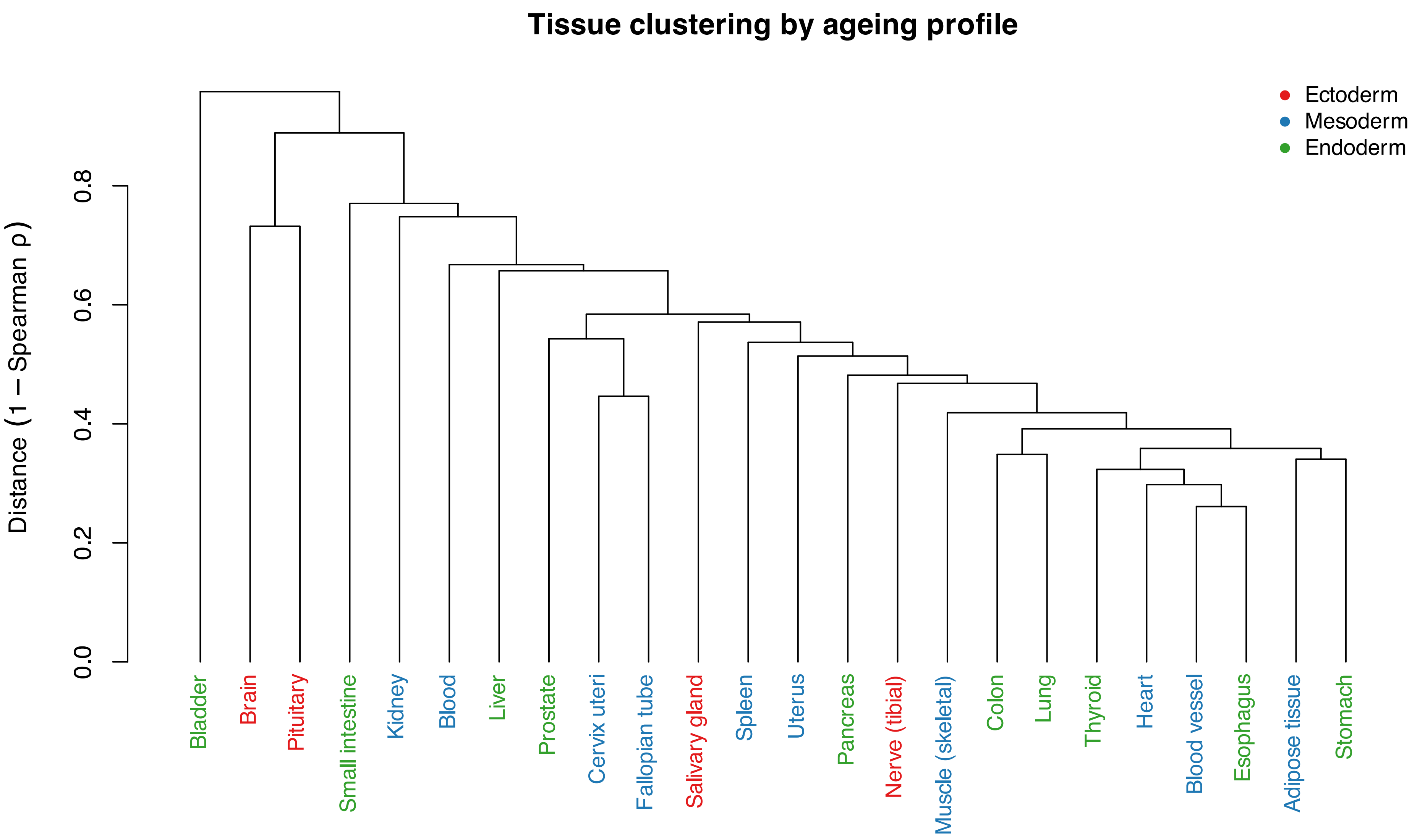
Tissue clustering by ageing profile
Distance (1 − Spearman ρ)
0.0
0.2
0.4
0.6
0.8
Ectoderm
Mesoderm
Endoderm
Bladder
Brain
Pituitary
Small intestine
Kidney
Blood
Liver
Prostate
Cervix uteri
Fallopian tube
Salivary gland
Spleen
Uterus
Pancreas
Nerve (tibial)
Muscle (skeletal)
Colon
Lung
Thyroid
Heart
Blood vessel
Esophagus
Adipose tissue
Stomach
